# Addition of adjuvant to DTaP modulates vaccine-induced immunological responses but is insufficient to improve protection in CD-1 mice

**DOI:** 10.1101/2023.03.03.530983

**Authors:** Kelly L. Weaver, Gage M. Pyles, Spencer R. Dublin, Annalisa B. Huckaby, Sarah J. Miller, Maria De La Paz Gutierrez, Emel Sen-Kilic, William T. Witt, Dylan T. Boehm, F. Heath Damron, Mariette Barbier

## Abstract

Pertussis is a vaccine-preventable respiratory disease caused by the Gram-negative bacterium *Bordetella pertussis*. While vaccination rates remain high in developed countries, incidence of pertussis has increased following the transition from wP vaccines to aP vaccines. The reemergence of pertussis is attributed, in part, to waning immunity induced by aP vaccination. Therefore, the objective of this work was to determine if addition of adjuvant to DTaP can modulate the immune response and improve protection compared to DTaP alone. In this study we immunized outbred, female CD-1 mice with 1/320^th^ the human dose of vehicle control, DTaP, and DTaP supplemented with adjuvant. Markers of early vaccine-induced memory were measured using a chemokine assay or by flow cytometry. Protection was assessed by measuring serological responses and quantifying bacterial burden in the respiratory tract at day 3 post-challenge. From this work we identified a partially protective aP vaccine dose to use for vaccination and challenge studies. We observed that MPLA and SWE promote robust anti-*B. pertussis* antibody responses and stimulate significant increases in early markers of vaccine-induced memory such as CXCL13, FDCs, and T_FH_ cells. Quil-A induced Th1 responses compared to DTaP alone, but none of the adjuvants improved protection against challenge with *B. pertussis*. Overall, the data suggests that addition of adjuvant modulates the protective immune responses induced by aPs. Further studies are needed to evaluate the B cell compartment and longevity of protection.

## 3.2 Introduction

Pertussis, otherwise known as whooping cough, is a vaccine preventable human respiratory disease^31^. Vaccines against pertussis have been implemented since the 1940’s and provide protection from severe disease and in the case of neonates and infants, death. While current pertussis vaccines have an excellent safety profile and protect against severe pertussis, they do not prevent transmission of pertussis^41,42^. Recent studies have shed also light on the waning immunity associated with acellular pertussis vaccination in which the vaccine-induced memory does not last long-term compared to whole cell vaccination. This insufficient protection raises the need for the development of next generation pertussis vaccines that can prevent both transmission and promote long-term protection.

It has long been hypothesized that addition of adjuvant to pertussis vaccines could improve the longevity of protection and pertussis vaccine-induced memory. Indeed, several groups have shown that addition of adjuvant improves protection and serological responses against *B. pertussis*^38,85,120,245,246^. However, there is still a lack in understanding of pertussis vaccine-induced memory responses. An adjuvant is a substance added to purified antigen-based vaccines, such as DTaP, to promote the immunogenicity. Adjuvants can broaden, alter, and increase immune responses^82,245^ by recruiting antigen presenting cells to the site of vaccination and aiding in the development of the adaptive immune responses. The mechanisms by which adjuvants work vary from depot effect to recruitment of innate immune cells, and the production of chemokines and cytokines^246,247^. By engaging these components, adjuvants can alter the adaptive immune response. Understanding how adjuvants affect the quantity and quality of adaptive responses allows for proper selection for vaccine development efforts.

There are a variety of substances that can act as adjuvants, such as salts, emulsions, saponins, and even microbial components, but few are licensed for use in humans^83,248^. The objective of this study was to determine if protection and vaccine-induced memory could be enhanced via evaluation of the immunopotentiation effects of each adjuvant when added to DTaP. The acellular pertussis vaccine DTaP is adjuvanted with aluminium hydroxide, or alum. The adjuvants selected include monophosphoryl A (MPLA), Quil-A, SWE, oligonucleotide (ODN) 1585, ODN 1826, and ODN 2395. Traditionally, alum or aluminium hydroxide has been utilized as a vaccine adjuvant. Aluminium salts commonly referred to as alum have been utilized in studies since 1926 and are thought to work via depot effect although mechanisms of action are not well understood. MPLA is a TLR4 agonist and has been shown to lead to a significant increase in T_FH_ cells, germinal center B cells, and plasma cells^249^. Quil-A is a saponin that stimulates cell mediated responses and antibody mediated responses^83,250^. SWE is an oil-in-water emulsion similar to MF59 and has been implemented in several preclinical trials^251,252^. MF59 induces potent T_FH_ cell responses and enhances germinal center B cell responses^253,254^. Finally, the last class of adjuvant evaluated in this study belongs to the TLR9 agonist group known as ODN’s that induce Th1 responses overall, but can have differing immunostimulatory effects^255–257^. Type A induces plasmacytoid dendritic cells (pDCs), B induces strong activation of B cells, and C induces both. It is for these reasons that the above adjuvants were chosen for our study.

In this work, we supplemented DTaP that is adjuvanted with alum with additional adjuvants. We hypothesized that the combination of adjuvants may function similarly to adjuvant systems which are combinations of immunostimulatory substances that can provide broad protection compared to a single adjuvant formulation. We assessed cell populations and chemokine responses associated with the development of immunological memory to understand how addition of adjuvant effects the immune response compared to both DTaP alone. These data suggest that while addition of adjuvant is not sufficient to improve protection against *B. pertussis* compared to DTaP alone, adjuvants have the potential to alter the immune response even in the presence of alum. We observed that Quil-A skewed the immune response from Th2 toward Th1, and MPLA and SWE induced significant increases in early markers of vaccine-induced memory such as CXCL13, FDC’s, and T_FH_ cells. Further studies are needed to analyze the B cell compartment and longevity of protection induced by addition of adjuvant to DTaP.

## 3.3 Materials and Methods

### 3.3.1 *B. pertussis* strains and growth conditions

*Bordetella pertussis* strain UT25Sm1 was kindly provided by Dr. Sandra Armstrong (University of Minnesota)^129,130^. UT25Sm1 strain has been fully genome sequenced (NCBI Reference Sequence: NZ_CP015771.1). UT25Sm1 was grown on Difco^™^ Bordet Gengou (BG) agar (VWR^™^, Cat. #90003-414) supplemented with 15% defibrillated sheep blood (Hemostat Laboratories, Cat. #DSB500) and streptomycin 100 μg/mL (Gibco^™^, Cat. #11860038) at 36°C for 48 hours. The number of viable bacteria and the Bvg+ (hemolytic and characteristic colony morphology) phenotype was confirmed to ensure consistency between each challenge. Bacteria were then collected using polyester swabs and resuspended in Stainer Scholte media^132^ (SSM) supplemented with L-proline and SSM supplement. SSM liquid culture was incubated for 24 hours at 36°C with constant shaking at 180 rpm until reaching mid-log phase OD_600nm_ 0.5 with 1 cm path width (Beckman Coulter^™^ DU 530 UV Vis spectrophotometer). The UT25Sm1 *B. pertussis* culture was diluted in supplemented SSM to OD_600nm_ = 0.24 - 0.245 (equivalent to 10^9^ CFU/mL) to be used for challenge or serological analysis by ELISA.

### 3.3.2 Vaccine preparation and immunization, bacterial challenge, and euthanasia

The acellular *B. pertussis* vaccine DTaP (Infranrix®, GlaxoSmithKline) was purchased. The following adjuvants were utilized in these studies: MPLA-SM VacciGrade<sup>™</sup> (InvivoGen, Cat. #vac-mpla), ODN 1585 VacciGrade<sup>™</sup> (InvivoGen, Cat. #vac-1581-1), ODN 1826 VacciGrade<sup>™</sup> (InvivoGen, Cat. #vac-1826-1), ODN 2395 VacciGrade<sup>™</sup> (InvivoGen, Cat. #vac-2395-1), Quil-A® adjuvant (InvivoGen, Cat. #vac-quil), and Sepivac SWE<sup>™</sup>, Seppic. SWE was mixed with 1/320^th^ the human dose of DTaP 1:1 bedside in the dark. MPLA, ODNs, and Quil-A were added to 1/320^th^ the human dose of DTaP at a concentration of 20 μg per mouse. All vaccines were intramuscularly administered at 1/320^th^ the human dose in 50 μL. The vaccines were diluted using endotoxin free phosphate buffered saline (PBS) (Millipore Sigma^™^, Cat. #TMS012A). PBS was administered as a vehicle control. In all experimental groups, 6-week-old outbred female CD1 mice (Charles River, Strain code 022) were used. Mice were primed at day 0, followed by a booster of the same vaccine at day 21. Non-challenged mice were euthanized at days 22 and 35 post-vaccination while challenged mice were only euthanized at day 35 post-vaccination. For challenged animals, mice were anesthetized three days before euthanasia by intraperitoneal injection (IP) ketamine (7.7 mg/kg) (Patterson Veterinary, Cat. #07-803-6637) and xylazine (0.77 mg/kg) (Patterson Veterinary, Cat. #07-808-1939) in sterile 0.9% NaCl (Baxter, Cat. #2F7124) and challenged intranasally with ∼2×10^7^ CFU/dose of live *B. pertussis* (10 µL per nostril)^120,191^. At day three post-challenge mice were euthanized by IP injection of Euthasol (390 mg pentobarbital/kg) (Patterson Veterinary, Cat. #07-805-9296) in sterile 0.9% *w/v* NaCl.

### 3.3.3 Quantification of bacterial burden

Lung and trachea homogenates as well as nasal lavage (nasal wash) were collected post mortem and used to enumerate bacterial burden per tissue. Mice were challenged at day 32 post-prime and processed three days later (day 35 post-vaccination). The nasal cavity was flushed with 1 mL sterile PBS for nasal lavage. The lung and trachea were homogenized separately in 1 mL sterile PBS using a Polytron PT 2500 E homogenizer (Kinematica). Samples were serially diluted in ten-fold dilutions in PBS and plated on BG agar to quantify viable bacterial burden. Plates were incubated at 36°C for 48-72 hours to determine colony forming units (CFUs) per mL.

### 3.3.4 Serological analysis of immunized mice

Enzyme linked immunosorbent assay (ELISA) was utilized to measure *Bordetella pertussis*-specific antibodies in the serum of immunized mice^191–193^. After euthanasia, blood was collected in BD Microtainer serum separator tubes (BD, Cat. #365967) via cardiac puncture at days 22 and 35 post primary immunization. Blood was centrifuged at 14,000 *x g* for 2 minutes and sera were stored at -80°C. Pierce^™^ high-binding 96 well plates (Thermo Scientific^™^, Cat. #15041) were coated with 5×10^7^ CFU/well viable *B. pertussis* overnight at 4°C. Plates were washed three times with PBS-Tween^®^20 (Fisher Scientific, Cat. #BP337-500), then blocked with 5% *w/v* non-fat dry milk (Nestle Carnation, Cat. #000500002292840) in PBS-Tween^®^20. Serum samples were serially diluted from 1:50 to 1:819,200 using 5% *w/v* milk in PBS-Tween^®^20. Plates were incubated at 37°C for 2 hours and washed four times with PBS-Tween^®^20. Secondary goat anti-mouse IgG antibody 1:2000 (Southern Biotech, Cat. #1030-04) conjugated to alkaline phosphatase was added and incubated for 1 hour at 37°C. Wells were washed five times with PBS-Tween^®^20 and Pierce p-Nitrophenyl Phosphate (PNPP) (Thermo Scientific, Cat. #37620) was added to each well to develop plates for 30 minutes in the dark at room temperature. The absorbance at 405 nm was read utilizing a SpectraMax^®^ i3 plate reader (Molecular Devices). The lower limit of detection for serum titers was 1:50, and for statistical analysis, all values below the limit of detection are represented with the arbitrary value of one. Endpoint titers were determined by selecting the dilution at which the absorbance was greater than or equal to twice that of the negative control.

### 3.3.5 Chemokine assay

CXCL13 levels were measured in sera from mice using the Mouse Magnetic Luminex® Assays (R&D Systems, Cat. #LXSAMSM) kit. Data was obtained using a Magpix (Luminex) instrument.

### 3.3.6 Tissue isolation, preparation, staining, and flow cytometry

Flow cytometry was used to characterize cell populations from the inguinal lymph nodes. Organs were harvested at days 22 and 35 post-prime. Lymph nodes were homogenized using disposable pestles (USA Scientific, Cat. #1405-4390) in Dulbecco’s Modified Eagle Media (DMEM) (Corning Incorporated, Cat. #10-013-CV) with 10% *v/v* fetal bovine serum (FBS) (Gemini Bio, Cat. #100-500). Homogenized samples were strained for separation using 70 μM pore nylon mesh (Elko Filtering Co, Cat. #03-70/33) and centrifuged for 5 minutes at 1,000 *x g*. Next, samples where resuspended in PBS with 5mM ethylenediaminetetraacetic acid (EDTA) (Fisher Scientific, Cat. #50-103-5745) and 1% *v/v* FBS. Single cell suspensions were incubated with 5 μg/mL anti-mouse CD16/CD32 Fc block (clone 2.4G2, Thermo Fisher Scientific, Cat. #553142) for 15 minutes at 4°C per the manufacturer’s instructions. Cells were stained with antibodies against cell surface markers (**Supplemental Table 1**) (FDCs:CD45^-^CD21/35^+^). Each single cell suspension was incubated with the antibody cocktail for 1 hour at 4°C in the dark. Samples were washed by resuspending in PBS, centrifuging, removing the supernatant, and washing in PBS with 5mM EDTA and 1% *v/v* FBS and fixed with 0.4% *w/v* paraformaldehyde (Santa Cruz Biotechnology, Cat. #sc-281692) overnight. After fixation, samples were centrifuged and washed before resuspension in PBS with 5mM EDTA and 1% *v/v* FBS. The samples were processed using an LSR Fortessa flow cytometer (BD Biosciences) and analyzed using FlowJo (FlowJo^™^ Software Version v10). Cells were counted using Sphero AccuCount 5-5.9 μm beads according to the manufacturer’s protocol (Spherotech, Cat. #ACBP-50-10).

### 3.3.10 ELISpot preparation and analysis

The Mouse IgG ELISpot (ImmunoSpot®, Cat. #mIgG-SCE-1M) was used to quantify *B. pertussis* specific antibody secreting cells in the bone marrow of immunized mice. UT25Sm1 was cultured as described above. PVDF membrane 96-well plates were coated with 5×10^7^ CFU/well *B. pertussis* and incubated overnight at 4°C. Bone marrow samples were isolated by centrifuging femurs at 400 *x g* for 5 minutes in 200 µL PCR tubes with holes in the bottom that were placed into 2 mL Eppendorf tubes. The bone marrow was resuspended in heat-inactivated filter-sterilized FBS and filtered through 70 µm mesh with FBS with 10% *v/v* dimethyl sulfoxide (DMSO) (Sigma-Aldrich, Cat. #D8418-100ML) and stored at -80°C. Bone marrow cells were thawed in a 37°C water bath and placed in DMEM with 10% *v/v* FBS. Cells were centrifuged at 400 *x g* for 5 minutes, resuspended in CTL Test B Culture medium (ImmunoSpot), diluted 1:10 with PBS and 1:1 with trypan blue stain (Invitrogen^™^, Cat. #T10282), and counted on the Countess II Automated Cell Counter (Invitrogen). Plates were washed with PBS and cells were added to the first row then serially diluted two-fold down the plate. Cells were incubated at 36°C overnight and imaged using the ImmunoSpot® S6 ENTRY Analyzer and CTL Software. Dilutions with spots ranging from ∼10-100 per well were selected to enumerate the number of anti-*B. pertussis* antibody-producing cells per sample. Cell counts were normalized to spots per 10^6^ cells using the cell and spot counts.

### 3.3.11 Statistics

Statistical analysis was performed using GraphPad Prism version 8 (GraphPad). When comparing three or more groups of parametric data a one-way ANOVA (analysis of variance) with Tukey’s multiple comparison test was used unless otherwise noted. For non-parametric data a Kruskal-Wallis test with Dunnet’s post-hoc test was used. The ROUT method was used to identify outliers when appropriate.

### 3.3.12 Animal care and use

All mouse experiments were approved by the West Virginia University Institutional Animal Care and Use Committees (WVU-AUCU protocol 1901021039) and completed in strict accordance of the National Institutes of Health Guide for the care and use of laboratory animals. All work was done using universal precautions at BSL2 under the IBC protocol # 17-11-01.

## 3.4 Results

### 3.4.1 Determining a partially protective vaccine dose

Previous studies from our laboratories have examined the differences in the immunological response to vaccination with aP and wP vaccines in the murine mouse model^258^. To do so, we have used highly saturating doses (1/10^th^ of the human dose) to be able to study longevity of protection and detect rare antigen-specific memory B cell populations. In more recent studies, we have however shown that it is important to use a non-saturating dose to determine the effect of antigen and adjuvant modification on the immune response to *B. pertussis* vaccination^119,255^. To determine a non-saturating dose of vaccine that would provide partial protection, we titrated the aP vaccine from 1/20th to 1/320th the human dose (**Figure 1**). Mice were boosted with the same vaccine at day 21 post-vaccination and challenged with 2×10^7^ CFUs/dose of *B. pertussis* at day 35 post-prime. Three days later, bacterial burden was quantified in the lungs of challenged mice. Blood was collected via cardiac puncture and used to measure the anti-PT antibody titers. We observed that all doses selected provided significant protection against bacterial colonization in the lung three days post-challenge compared to the mock-vaccinated mice (**Figure 1**), and that protection was dose-dependent. The 1/20^th^ dose provided the highest protection, with a bacterial burden in the lung three days post-challenge lower than the limit of detection. We also observed a dose-dependent production of anti-pertussis antibodies in response to vaccination, with the highest titers observed in mice vaccinated with 1/20^th^ of the human dose and the lowest titers in the mice vaccinated with 1/320^th^ of the human dose (**Figure 1**). The dose 1/320th was selected to move forward as it conferred the least protection amongst immunized mice with the exception of the PBS challenged group.

**Figure 1.**
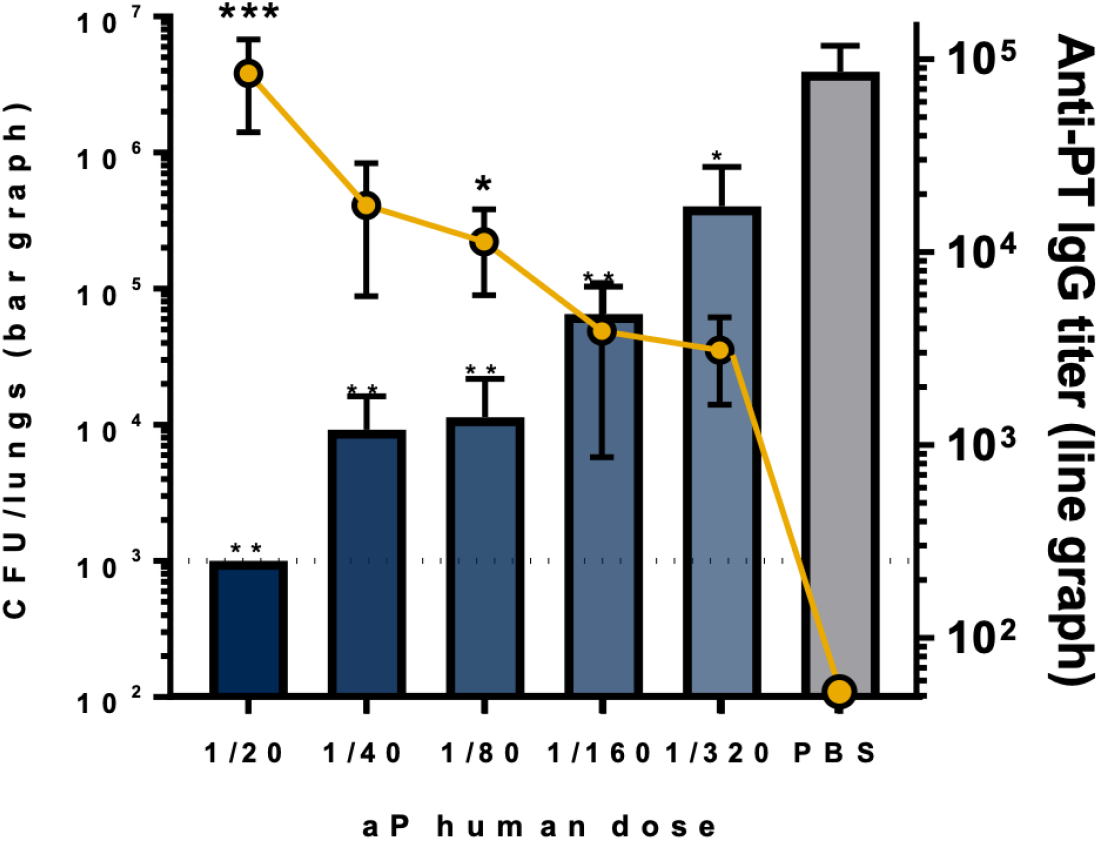
Selection of a partially protective vaccine dose. Anti-PT IgG titers measured in the sera of immunized mice compared to bacterial burden quantified in the lungs. The p-values were calculated for each time point using ANOVA followed by a Tukey’s multiple-comparison test, ^*^*p* < 0.05, ^**^*p* < 0.01, and ^***^*p* < 0.001. Error bars are mean ± SEM values.

### 3.4.2 MPLA and SWE promote robust anti-*B. pertussis* antibody responses

One of the primary correlates of protection measured to determine pertussis vaccine efficacy is the antibody response to the pathogen. To determine if modification of the adjuvant content of the DTaP vaccine modifies the serological response, we first performed immunogenicity studies in which 1/320th the human dose of vehicle control, DTaP only, or DTaP+adjuvant was administered intramuscularly to mice on days 0 and 21. In this first phase of the study, the adjuvants Quil-A, MPLA, SWE, ODN 1585, ODN 1826, and ODN 2395 were evaluated. At day 32 mice were intranasally challenged with 2×10^7^ CFU of *B. pertussis* and bacterial burden was quantified in the respiratory tract from the lungs, trachea, and nasal wash (**Figure 2A**). From the serum anti-*B. pertussis* IgG antibodies were measured by ELISA (**Figure 2B**). All mice vaccinated with DTaP alone or combined with various adjuvants produced detectable levels of anti-*B. pertussis* antibody levels. However, we observed **A** a statistically significant increase in anti-*B. pertussis* antibody levels in mice vaccinated with DTaP+MPLA or DTaP+SWE compared to non-vaccinated mice. These data suggest that addition of MPLA and SWE to DTaP could potentiate the antibody responses against *B. pertussis* compared to alum alone contained in DTaP. These two adjuvants were therefore selected for subsequent characterization of the immune response they generated. In addition, as addition of Quil-A negatively affected antibody production to *B. pertussis*, this adjuvant was also selected for further characterization to determine its effect on the immunological response to DTaP.

**Figure 2.**
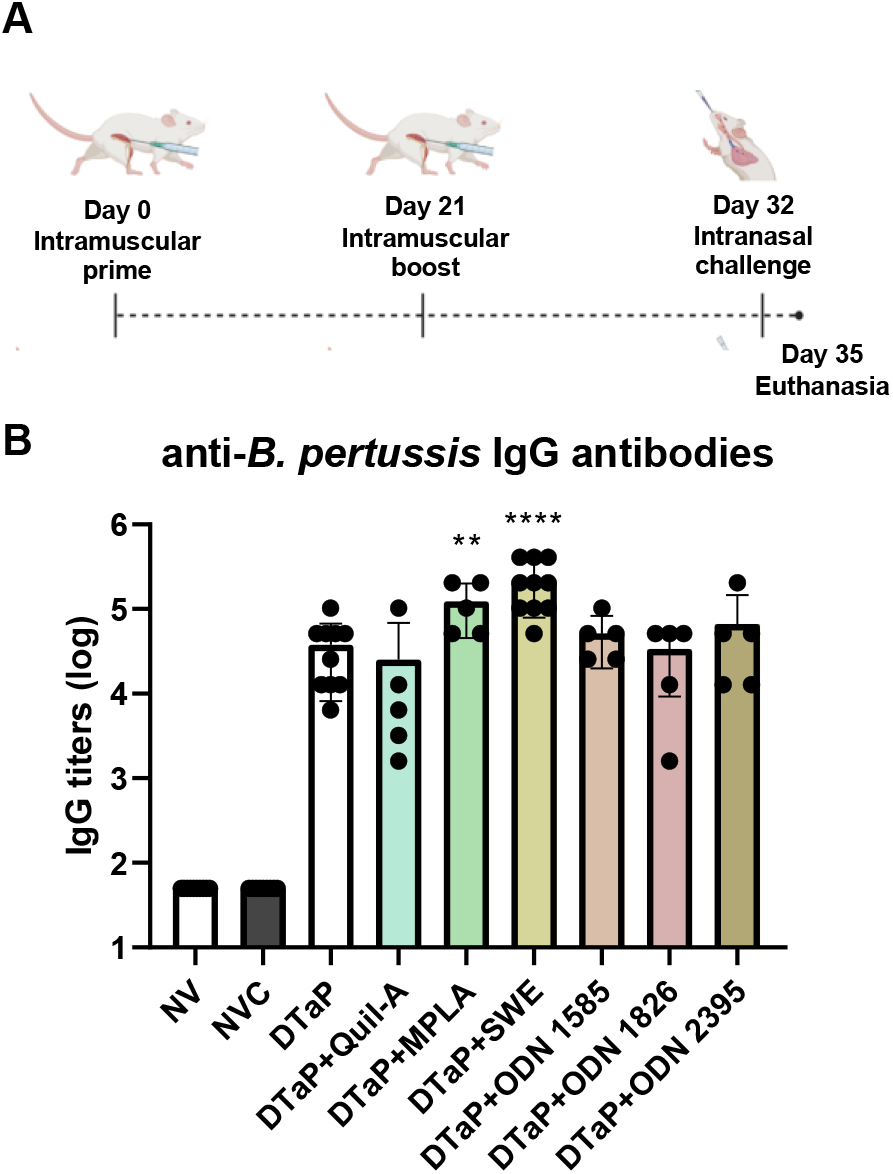
MPLA and SWE promote robust anti-*B. pertussis* antibody responses. **A)** Experimental design. Mice were vaccinated on days 0 and 21 with 1/320th the human dose of each vaccine. On day 32 mice were intranasally challenged with 2×10^7^ CFU/ dose of B. pertussis, and euthanized three days later. **B)** Serology showing anti-B. pertussis IgG antibody responses. The p-values were calculated for each time point using ANOVA followed by a Tukey’s multiple-comparison test, ^**^*p* < 0.01, and ^****^*p* < 0.0001. Error bars are mean ± SEM values (n=5-10 per group).

### 3.4.3 Addition of Quil-A to DTaP alters the type of Th cell response compared to DTaP alone

aP vaccines are known to induce Th2 responses, however, some of the adjuvants selected are known to drive Th1 responses. Therefore, we next assessed if addition of adjuvant could skew the Th2 response as measured by analyzing the anti-*B. pertussis* IgG1/IgG2a ratio (**Figure 3**). The IgG1/IgG2 ratio was measured from the sera of immunized mice. The only significant difference observed compared to DTaP was in the Quil-A adjuvanted group in which we observed a shift toward a Th1 response. Addition of MPLA and SWE did not significantly alter the humoral response compared to DTaP alone and both vaccines were associated with a Th2-dominant response. Interestingly, some of the mice vaccinated with DTaP + MPLA had a very low IgG1/IgG2a ratio, suggesting that they developed a different Th response more skewed towards Th1 compared to the mice administered DTaP alone. Overall, the data suggest that addition of adjuvant to DTaP has the potential to skew the type of protective immune response from a Th2-dominant response to a more balanced Th1-Th2 response.

**Figure 3.**
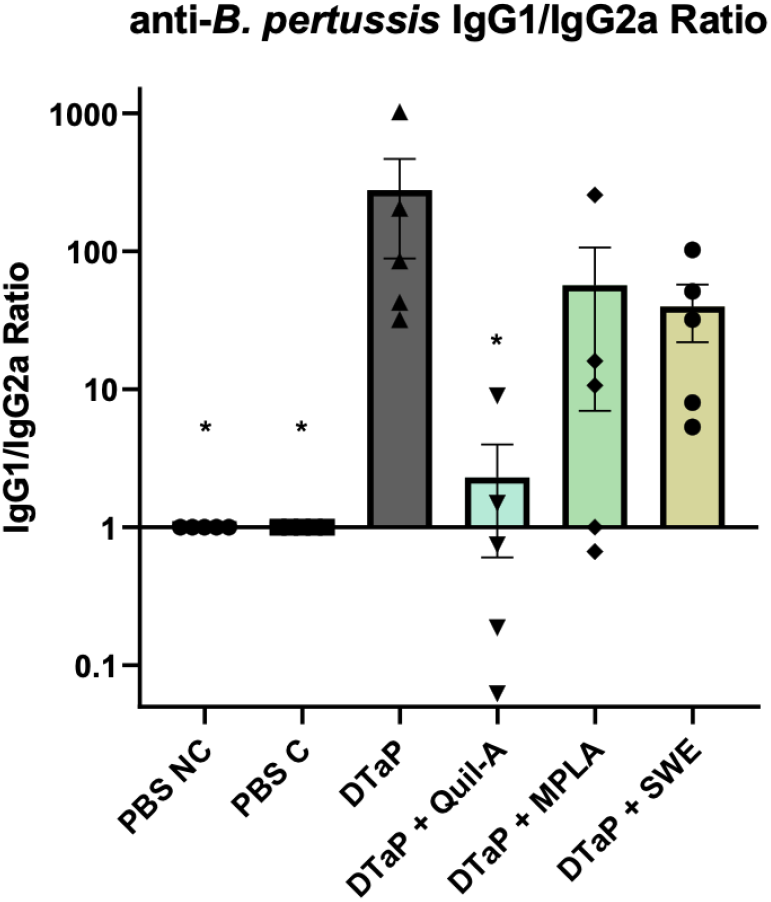
Addition of Quil-A induces Th1 responses compared to DTaP alone. The anti-*B. pertussis* IgG1/IgG2a ratio was measured in the sera of immunized mice. The p-values were calculated for each time point using ANOVA followed by a Tukey’s multiple-comparison test, ^*^*p* < 0.05. Error bars are mean ± SEM values (n=5 per group).

### 3.4.4 SWE and MPLA potentiate increases in early markers of vaccine-induced memory compared to DTaP alone

To determine if addition of adjuvant impacts pertussis vaccine-induced memory, early markers of germinal center formation were measured in non-challenged mice (**Figure 4**). CXCL13 levels were measured in blood collected via submandibular bleeding at days 1, 7, 14, 20, and 28 following vaccination.

**Figure 4.**
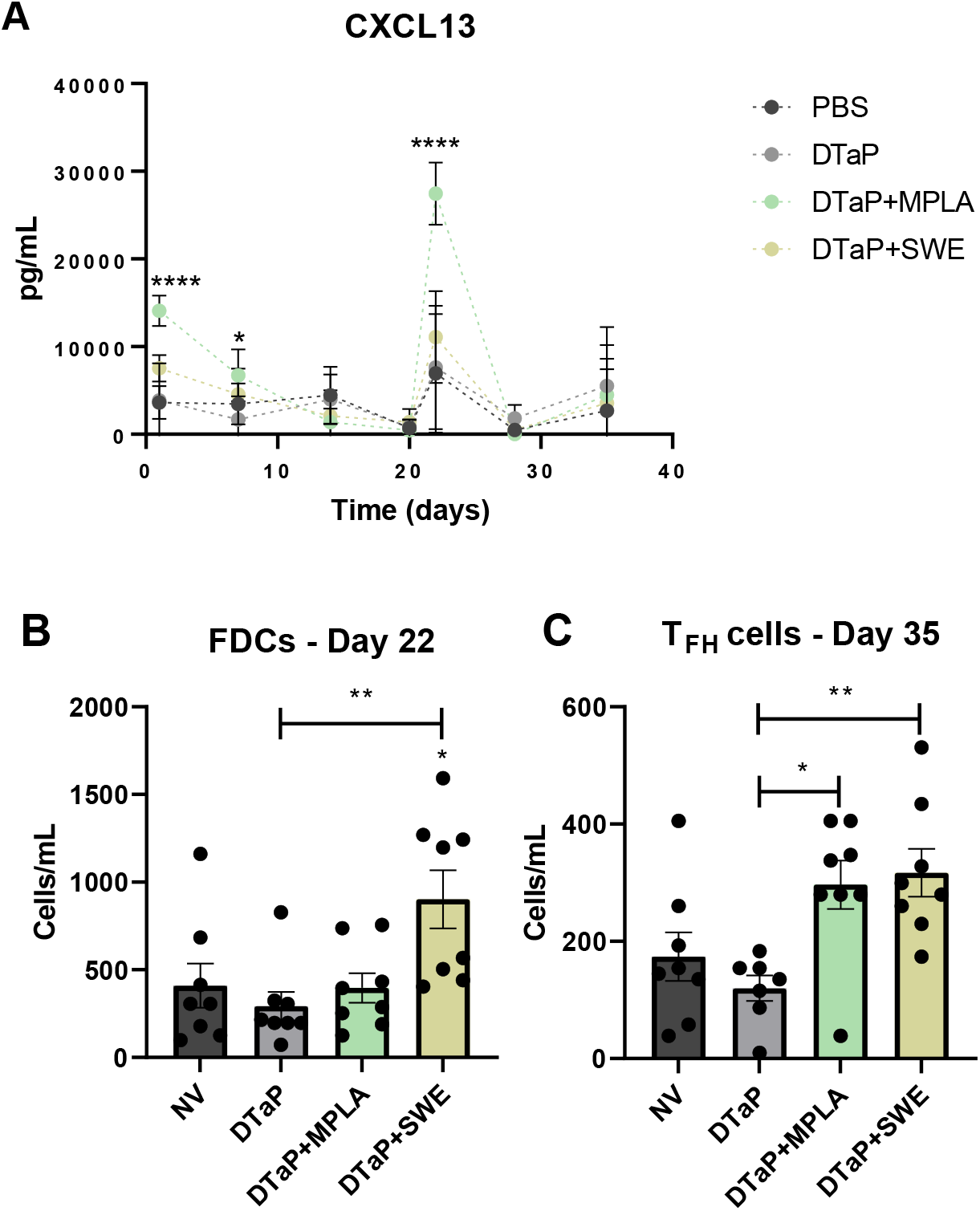
SWE and MPLA potentiate increases in early markers of vaccine-induced memory compared to DTaP alone. A) CXCL13 levels measured in the sera of non-challenged CD-1 mice weekly following vaccination and boost. B) Follicular dendritic cells measured at day 22 following prime in the draining lymph node of immunized mice. C) T follicular helper cells at day 35 post-vaccination in the draining lymph node of immunized mice. The p-values were calculated for each time point using ANOVA followed by a Tukey’s multiple-comparison test, ^*^*p* < 0.05 and ^****^*p* < 0.0001. Error bars are mean ± SEM values (n=10 per group).

Blood samples were collected by cardiac puncture at days 22 and 35 as secondary method of euthanasia. A significant increase in CXCL13 was only observed at day 22 post-prime in DTaP+MPLA vaccinated mice compared to DTaP only (**Figure 4A**). This data suggests that addition of MPLA to DTaP leads to an increase in CXCL13 and potentially long-term memory as CXCL13 is a biomarker of germinal center formation. FDCs present antigen via complement receptors and play a critical role in the differentiation of T_FH_ cells and the development of germinal centers. At day 22 post-prime, we measured FDCs from the draining lymph nodes of immunized mice. We observed a significant increase in FDCs elicited by addition of adjuvant to DTaP compared to DTaP alone (**Figure 4B**). By day 35 post-prime T_FH_ levels were significantly increased in the presence of both MPLA and SWE compared to DTaP alone (**Figure 4C**). Altogether, these data suggest that the addition of both MPLA and SWE to DTaP has the potential to alter early pertussis vaccine-induced memory responses.

### 3.4.5 SWE induces a significant increase in *B. pertussis* specific antibody secreting cells compared to DTaP alone

Antibodies play a crucial role in vaccine-mediated protection via neutralization, opsonization, and activation of complement^259^. High affinity antibodies are produced by antibody secreting cells that differentiate from naive B cells as a result of germinal center activity. To evaluate if addition of adjuvant impacts B cell responses, *B. pertussis* specific ASCs were measured in the bone marrow of non-challenged CD-1 mice at day 35 post-vaccination (**Figure 5**). We observed that the addition of SWE to DTaP led to a statistically significant increase in *B. pertussis*^+^ ASCs compared to both vehicle control and DTaP alone. MPLA did not lead to an increase in *B. pertussis*^+^ ASCs in comparison to vehicle control nor DTaP alone. These data suggest that not only does SWE have the potential to increase early markers of germinal center formation, but it also influences the B cell compartment.

**Figure 5.**
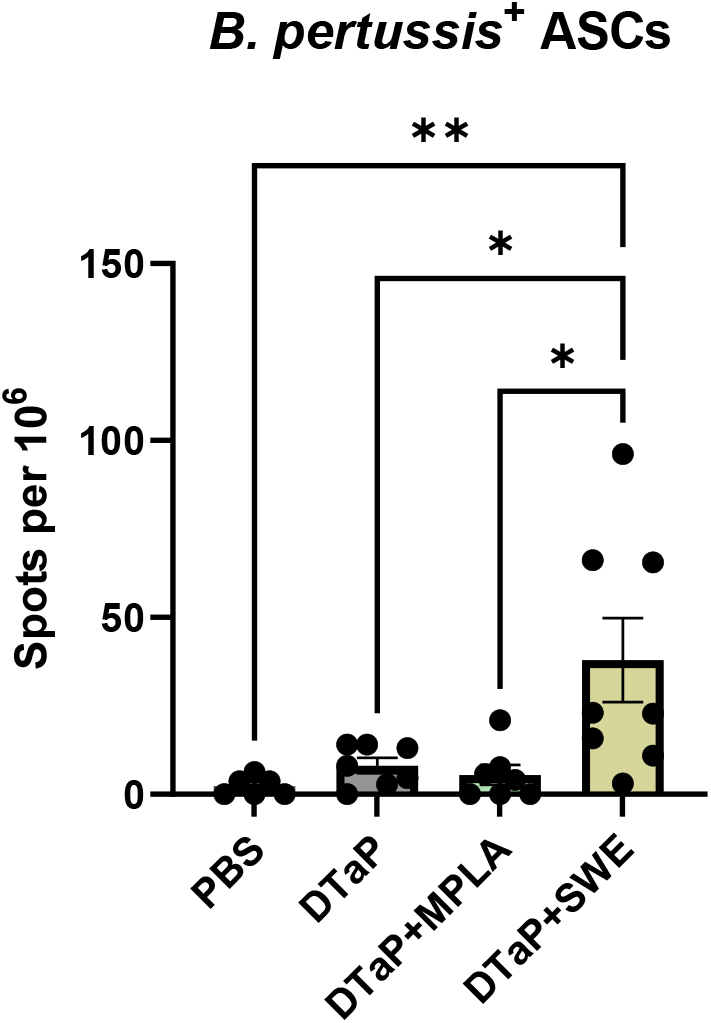
SWE induces a significant increase in *B. pertussis* specific antibody secreting cells compared to DTaP alone. ASCs were measured from the bone marrow of immunized mice at day 35 post-vaccination by ELISpot. ROUT was used to identify outliers and the *p*-values were calculated using ANOVA followed by a Tukey’s multiple-comparison test, ^*^*p* < 0.05 and ^**^*p* < 0.01. Error bars are mean ± SEM values (n=10 per group).

### 3.4.6 Adjuvant modification is insufficient to improve protection against challenge with *B. pertussis*

Antibodies produced in response to DTaP vaccination play a role in bacterial opsonization and in neutralizing the activity of pertussis toxin, one of the major virulence factors of *B. pertussis*. During challenge with *B. pertussis*, pertussis toxin is released, leading to an increase in circulating neutrophils called leukocytosis. Neutralization of pertussis toxin activity by circulating antibodies can be measured as a decrease in leukocytosis in mice after challenge. To determine if the increase in *B. pertussis* antibody production associated with addition of MPLA and SWE to the DTaP vaccine (**Figure 4B**) is associated with an increase in pertussis-toxin neutralizing and decrease in leukocytosis, the number of circulating neutrophils were measured from the whole blood of challenged female CD-1 mice three days post-challenge (**Figure 6A**). There was a significant decrease in neutrophil levels observed in all groups compared to the non-vaccinated challenged group. Surprisingly, addition of Quil-A to DTaP provided the least amount of protection against leukocytosis (1.484 K/µL). The lowest number of neutrophils was observed in the non-vaccinated and DTaP+SWE immunized group suggesting that SWE may be a promising candidate to evaluate further.

**Figure 6.**
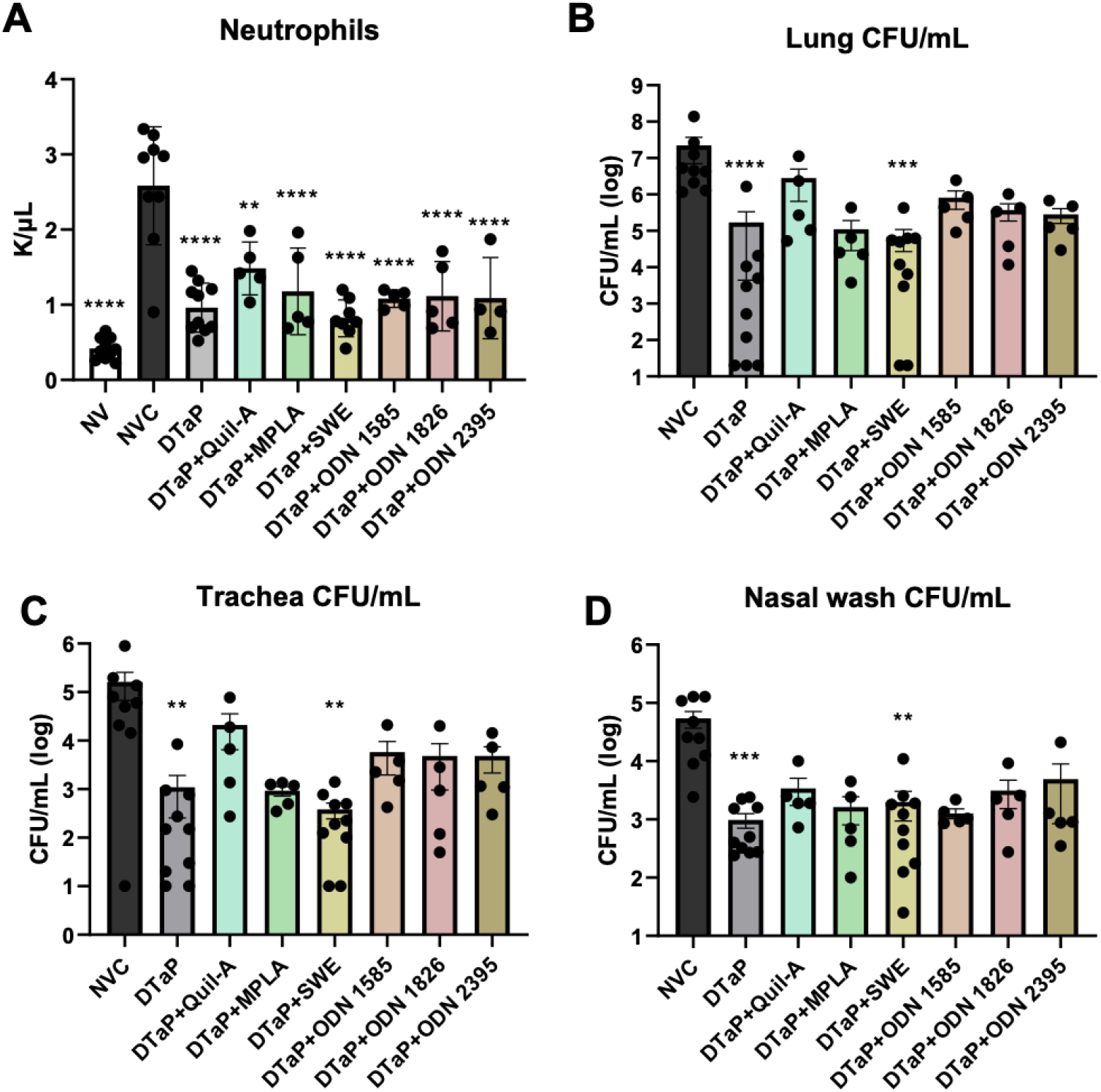
Adjuvant modification is insufficient to improve protection against challenge with *B. pertussis*. **A)** Neutrophils measured in whole blood of mice at day 3 post-challenge. **B-D)** Bacterial burden was measured in the **B)** lungs, **C)** trachea, and **D)** nasal wash of immunized mice via serial diluting and colony counting. The p-values were calculated for each time point using ANOVA followed by a Tukey’s multiple-comparison test, ^**^*p* < 0.01, ^***^*p* < 0.001, and ^****^*p* < 0.0001. Error bars are mean ± SEM values (n=5-10 per group).

Bacterial burden was measured in the lungs (**Figure 6B**), trachea (**Figure 6C**), and nasal wash (**Figure 6D**) of challenged mice. Data is represented as CFU/mL for each organ. A significant decrease in bacterial burden was observed in the DTaP and DTaP+SWE groups compared to non-vaccinated challenged mice in all organs. While there was no significant decrease in bacterial burden in the MPLA adjuvanted group, the trend was similar to the SWE adjuvanted group. The same trends were observed in the trachea and nasal wash of immunized mice, yet the most protection was observed in the lung. The data suggest that differences in adjuvanted groups compared to DTaP alone are not observed when measuring protection via quantification of bacterial burden.

These studies shed light on the opportunity for future formulation of DTaP vaccines with adjuvant systems to modulate the immune response to vaccination and improve protection conferred by aP vaccines. While there were no differences observed in protection, the data suggest that the addition of adjuvant can skew the type of immune response compared to DTaP alone. Addition of adjuvant to DTaP improved the early vaccine-induced memory responses such as CXCL13 levels, FDCs, and T_FH_ cells. Further studies need to be done to evaluate the B cell compartment and longevity of protection conferred by addition of MPLA and SWE compared to DTaP alone. Overall, these data provide evidence that next-generation pertussis vaccines could potentially benefit from the implementation of adjuvant systems.

## 3.5 Discussion

Current aPs exhibit an excellent safety profile, yet the protection conferred is suboptimal. aPs do not prevent transmission or colonization of the respiratory tract by *B. pertussis*, nor do they provide long-term protection^42,44^. Evidence shows that although vaccine coverage is high, overall incidence and outbreaks of pertussis have increased in DTaP primed individuals. Waning immunity, or a decrease in protection over time following vaccination, contributes to the suboptimal protection associated with aPs^38,160^. The mechanisms underlying aP induced protection are not well understood, and biomarkers of long-term protection are needed for future vaccine development.

In our studies we vaccinated mice with a non-saturating dose, 1/320th the human dose of DTaP, with or without addition of adjuvant (**Figure 2**). It was critical to use a partially protective vaccine dose to able to evaluate the effects of each adjuvant. Outbred CD-1 mice were primed at day 0 and boosted at day 21. Blood was collected via cardiac puncture and anti-*B. pertussis* IgG antibody titers were measured at day 35 post-vaccination. While not statistically increased compared to DTaP alone, MPLA and SWE stimulated robust antibody responses (**Figure 2B**). Although levels of anti-*B. pertussis* IgG antibody responses are not statistically improved, the functionality of antibodies induced by MPLA and SWE could differ from DTaP alone. Functionality of antibodies is critical to consider during vaccine development, especially in the case of pertussis. aP vaccines generate two main categories of antibodies, those that are bactericidal and those that mediate phagocytosis and clearance of the bacteria. In the DTaP vaccine antibodies generated against pertactin primarily mediate bactericidal responses while anti-filamentous hemagglutinin antibodies promote phagocytic uptake^260–262^. As over 85% of *B. pertussis* clinical isolates no longer express pertactin, it is also crucial to consider the need for new antigens that drive bactericidal antibodies against antigens on the surface of the bacterium^263^. Therefore, the functionality, affinity, and avidity of antibodies should be evaluated in future studies.

Next, the IgG1/IgG2a ratio was measured to determine if adjuvants could shift the T cell response from Th2 dominant observed with DTaP alone, toward Th1 (**Figure 3**). The data suggests that even in the presence of alum, the addition of adjuvant impacts T cell responses. Although a balance between Th1/Th2 responses could be beneficial, this does not indicate that protection will be improved. Quil-A potentiated a Th1 dominant phenotype, decreasing the Th2 bias, but did not lead to a statistically significant reduction in bacterial burden in the respiratory tract compared to NVC mice (**Figure 6B-D**). These data are important to consider given that T cell responses are a topic of debate in pertussis vaccine development efforts. wPs and natural infection are associated with the induction of Th1/Th17 dominant responses^48^. It is hypothesized that Th1/Th17 responses may contribute to the long-term protection observed in wP and convalescent individuals whereas aPs stimulate Th2 responses and confer insufficient, short-term protection.

Interestingly, MPLA and SWE altered early markers of vaccine-induced memory (**Figure 4**). MPLA led to a significant increase in CXCL13 levels as early as one day post-vaccination compared to DTaP alone (**Figure 4A**). CXCL13 in the MPLA group peaked immediately following boost at day 22 post-vaccination. As CXCL13 is a biomarker for germinal center formation, MPLA could potentially enhance the longevity of memory conferred by DTaP. Previous studies suggest that MPLA induces robust T_FH_ cell, germinal center B cell, and plasma cell responses leading to the production of neutralizing antibodies. While MPLA did not lead to a significant increase in FDCs at day 22 (**Figure 4B**), it did induce a significant increase in T_FH_ cells at day 35 compared to DTaP alone (**Figure 4C**). SWE did not promote significant increases in CXCL13, but did significantly increase FDCs and T_FH_ cells suggesting that it also has the potential to improve pertussis vaccine-induced memory. Next, *B. pertussis* specific ASCs were measured in the bone marrow of immunized mice at day 35 post-prime to assess the impact of adjuvant on the B cell compartment. We observed that SWE led to a significant increase in *B. pertussis*^+^ ASCs in the bone marrow (**Figure 5**). These data suggest that addition of adjuvant to DTaP influences different aspects of the vaccine-induced memory response. The mechanisms underlying these findings, as well as the longevity of memory need to be further evaluated.

Altogether, given these findings, these data suggest that adjuvants have immunopotentiating effects when added to DTaP that can skew T cell responses, improve vaccine-induced memory, but do not enhance protection. While Quil-A balanced T cell responses toward Th1/Th2 and significantly decreased leukocytosis (**Figure 6A**), it did not decrease bacterial burden compared to NVC mice. Similarly to Quil-A, none of the ODNs selected improved protection compared to NVC and were not evaluated further. MPLA and SWE led to robust antibody responses and protection from challenge and were selected for further evaluation. As MPLA and SWE improved early markers of vaccine-induced memory we propose moving forward to measure *B. pertussis* specific MBC and ASC over time to determine if the longevity of protection can be enhanced compared to DTaP alone. These data provide evidence for the exploration into adjuvant systems for the development of next-generation vaccines that can prevent transmission, colonization, and induction of long-term protection.

## 3.6 Acknowledgements

The authors would like to acknowledge Dr. Kathleen Brundage, director of the flow cytometry and single cell core at West Virginia University in Morgantown, WV.

## 3.7 Contributions

KLW, MB, and FHD were responsible for study design. KLW led experiments. GMP, SRD, ABH, SJM, ESK, MPG, WTW assisted in data collection. KLW, MB, and FHD were responsible for writing the manuscript and all authors approved and provided edits.

---

**Supplementary table 1.** T_FH_ and FDC flow cytometry panels.

| T Follicular Helper Cells |  |  |  |
| --- | --- | --- | --- |
| Antibody | Fluorophore | Company | Catalog Number |
| CD185 | PE | BD Biosciences | 551959 |
| PD-1 | Per-CP-eFluor710 | eBioscience | 46-9985-82 |
| CD4 | APC-Cy7 | BioLegend | 100526 |
| CD3ε | BV510 | BD Biosciences | 563024 |
| Follicular Dendritic Cells |  |  |  |
| Antibody | Fluorophore | Company | Catalog Number |
| CD45 | PE-CF594 | BD Biosciences | 562420 |
| CD21/CD35 | BV421 | BD Biosciences | 562756 |

